# Are we moving the dial? Canadian Health Research Funding Trends for Women’s Health, 2S/LGBTQ+ Health, Sex, or Gender Considerations

**DOI:** 10.1101/2022.11.30.518613

**Authors:** Tori N. Stranges, Amanda B. Namchuk, Tallinn F. L. Splinter, Katherine N. Moore, Liisa A. M. Galea

## Abstract

**Background:** Sex and gender impacts health outcomes and disease risk throughout life. The health of women and members of the Two-Spirit, Lesbian, Gay, Bisexual, Transgender, Queer or Questioning, Intersex, and Asexual (2S/LGBTQ+) community is often compromised as they experience delays in diagnosis. Distinct knowledge gaps in the health of these populations has prompted funding agencies to mandate incorporation of sex and gender into research. Sex-and gender-informed research perspectives and methodology increases rigor, promotes discovery, and expands the relevance of health research. Thus, the Canadian Institutes of Health Research (CIHR) implemented a Sex and Gender-based Analysis (SGBA) framework recommending the inclusion of SGBA in project proposals in 2010 and then mandating the incorporation of SGBA into grant proposals in 2019. To examine whether this mandate resulted in increased mention of sex or gender in funded research abstracts, we searched the publicly available database of grant abstracts funded by CIHR to analyze the percentage of abstracts that mentioned sex or gender of the population to be studied. To better understand broader health equity issues we also examined whether the funded grant abstracts mentioned either female-specific health research or research within the 2S/LGBTQ+ community.

**Results:** We categorized a total of 8,964 Project and Operating grant abstracts awarded from 2009- 2020 based on their study of female-specific or a 2S/LGBTQ+ populations or their mention of sex or gender. Overall, under 3% of grant abstracts funded by CIHR explicitly mentioned sex and/or gender, as 1.94% of grant abstracts mentioned sex, and 0.66% mentioned gender. As one of the goals of SGBA is to inform on health equity and understudied populations with respect to SGBA, we also found that 5.92% of grant abstracts mentioned female-specific outcomes, and 0.35% of grant abstracts focused on the 2S/LGBTQ+ community.

**Conclusions:** Although there was an increased number of funded grants with abstracts that mentioned sex and 2S/LGBTQ+ health across time, these increases were less than 2% between 2009 to 2020. The percentage of funded grants with abstracts mentioning female-specific health or gender differences did not change significantly over time. The percentage of funding dollars allocated to grants in which the abstracts mentioned sex or gender also did not change substantially from 2009-2020, with grant abstracts mentioning sex or female-specific research increasing by 1.26% and 3.47% respectively, funding allocated to research mentioning gender decreasing by 0.49% and no change for 2S/LGBTQ+-specific health. Our findings suggest more work needs to be done to ensure the public can evaluate what populations will be examined with the funded research with respect to sex and gender to advance awareness and health equity in research.

**Highlights:**

- The percentage of funded grants in which the abstracts mentioned sex or gender in health research remained largely unchanged from 2009 to 2020 with the largest increase of 1.57% for those mentioning sex.
- Total funding amounts for grants that mentioned sex or gender in the abstract stagnated or declined from 2009 to 2020.
- The percentage of funded grants in which the abstracts focusing on female-specific health did not change across 2009-2020, but the percentage of funding dollars increased by 3.47%.
- The percentage of grants in which the abstracts mentioned 2S/LGBTQ+-specific health more than tripled across 2009-2020 but remained less than 1% of all funded grants.

## Background

Sex and gender play significant roles in health outcomes and disease risk (1). Sex refers to the biological attributes of females, males and intersex individuals, whereas gender is a psychosocial construct based on gender identity and society’s expectations of roles and behaviors based on that construction. Examining the contribution of sex and gender in research is essential to our complete understanding of the mechanisms that contribute to the etiology, manifestation, and response to treatment of disease (1,2). The importance of inclusion cannot be overstated. For example, in humans genetic polymorphisms and differences in gene expression are often distinct between males and females (3,4), suggesting that any treatment could have disparate effects by sex. The European Gender Equality Index of 2021 highlights that access to power and resources based on sex, gender, and sexual orientation combined with intersecting social categories determines health outcomes (5). As health research in this area progresses, funding agencies must continue to incentivize and highlight the importance of sex and gender based analysis (SGBA) to reduce health disparities and improve health outcomes.

Health outcomes differ in a host of diseases and across the entire spectrum of medicine between males, females and intersex individuals. Sex differences in diseases exist in manifestation (6–9), diagnosis time (10), misdiagnosis (9), treatment efficacy (11), and progression of disease (10). For example, females demonstrate poorer outcomes and increased mortality due to cardiovascular disease compared to males (11). However, clinical trials in cardiovascular disease continue to recruit more males than females (12). Female deaths attributed to cardiovascular disease outnumber deaths by breast, ovarian, uterine, cervical, and vaginal cancers combined (13). Yet, diseases with the highest burden on females remain chronically underfunded, whereas diseases that afflict primarily males are more likely to be appropriately or overfunded relative to disease burden (14). Within-sex disparities exist as well, as breast cancer research receives an outsized proportion of funding compared to disease burden across females, whereas endometriosis receives proportionately fewer research dollars (14–16). Although attention to sex and gender is an important step towards health equity, in order to understand why disparities in health outcomes exist between sexes and gender identities, it is also imperative to conduct within-sex and gender comparisons as well. Female sex-specific experiences, such as menses, menopause, and pregnancy, impact health outcomes and disease risk of conditions that target every organ including the heart, lung, kidney, and brain (17–20) and studying these effects do not require a comparison to males. Thus, the lack of health research into both sex differences and female-specific factors perpetuates disparities in duration of diagnoses, and the side effects due to treatments– resulting in devastating health effects for females compared to males (10,11,21,22).

Many of these health discrepancies can be attributed to the lack of female data collection in both animal models and clinical trials in North America. Starting in the late 1970’s, all premenopausal females were banned from participating in drug trials for almost two decades by the Food and Drug Administration (FDA) in the United States due to concerns of causing harm to the reproductive process (22). The importance of considering sex and gender differences in research has prompted funding agencies, including the Canadian Institutes of Health Research (CIHR), to implement SGBA frameworks for grant applications and adjudication. Figure 1 shows a timeline of funding agencies’ changes in policy to encourage inclusion of sex and gender. In 1993, the National Institutes of Health (NIH) mandated the inclusion of women in clinical trials, but to this day the inclusion of women in clinical trials is only a recommendation in Canada. Female health and sex differences continue to be overlooked in both clinical trials and basic research as women and female health are still routinely underrepresented (22–24). In the United States, even in research that includes women and females, the majority of these investigations (80%) do not use sex or gender as a factor in their primary outcomes, nor do they ask questions on whether female-specific factors influence these outcomes (25). This practice negates the benefits of diverse representation in research and does little to close the gap in our understanding of how sex and gender impact health.

**Figure 1.**
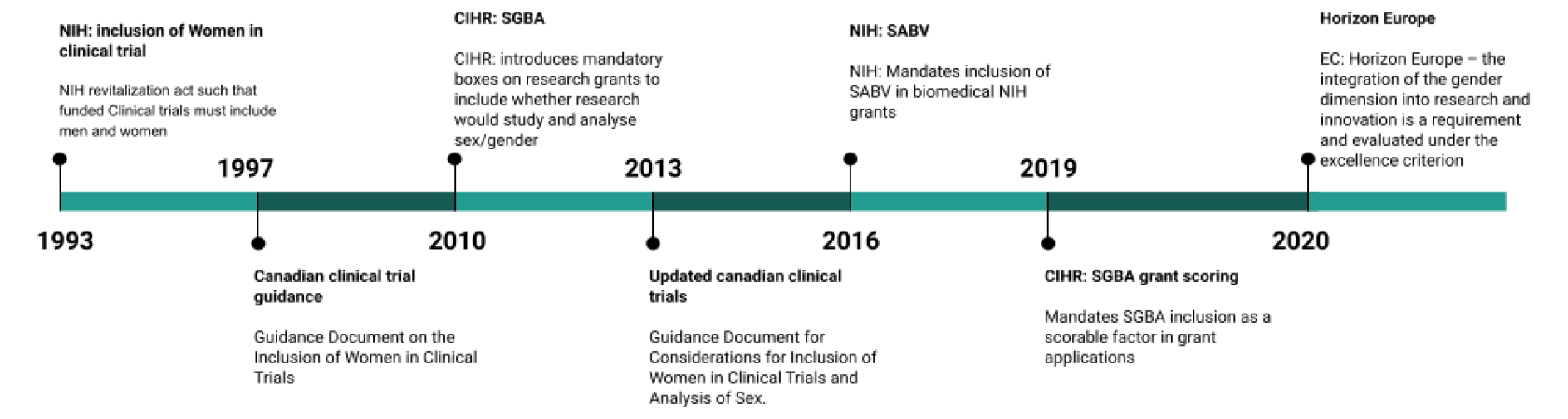
Timeline of mandates for inclusion and analysis of women/females in research from 1993 to 2020 from the National Institutes of Health (NIH) in the United States, the Canadian Institutes of Health Research (CIHR), and Horizon Europe. Prior to 1993 all premenopausal females were banned by the Food and Drug Administration (FDA) from participating in drug trials in the United States.

Studying sex and gender influences is only one step in understanding why health disparities exist across sex or gender. Gender identity and sexual orientation are also key considerations for health research from a health equity lens. The term Two-Spirited, Lesbian, Gay, Bisexual, Transgender, Queer or Questioning (2S/LGBTQ+) includes individuals who have both diverse gender identities as well as sexual orientations. Gender plays a role in healthcare seeking behavior, symptom perception by clinicians, disease diagnosis, and treatment (1,10,26,27). Sexual orientation is also a driving factor for poor health outcomes (28). 2S/LGBTQ+ community members often face discrimination and trauma based on their gender, sexual, and racial identities when accessing health and social services and experience poorer health outcomes as a result (28,29). Despite ample literature demonstrating that gender-affirming healthcare improves health outcomes and mitigates risk of stigma and discrimination (30–33), evidence continues to suggest that 2S/LGBTQ+ community members experience increased disease risk and poorer health outcomes (29,34,35).

Although more research is being conducted in males and females or considering gendered health experiences as well as sexual orientation, little headway has been made to increase the analysis or examination of potential sex and gender differences in published research (23,33,36–39). In the wake of the COVID-19 pandemic and subsequent vaccine trials, these problems persisted as sex and gender differences continued to be overlooked. Only 4% of COVID-19 clinical trials explicitly reported a plan to include sex or gender as an analytical variable (39). Further, during the initial COVID-19 vaccine trials, questions on menstrual, menopause disturbances or the effects on pregnancy were not specifically probed, likely leading to greater vaccine hesitancy among women (40). COVID-19 vaccine uptake is just one example of how neglecting sex, gender, and female-specific health in research can directly hinder public health efforts and reduce health equity.

Research funding bodies have acknowledged the lack of progress in these areas. The European Commission reported a mere 3% increase in grant applications with a gender dimension from 2015-2017 (41) and the NIH has acknowledged that the improved inclusion of women in clinical trials has not done enough to improve Sex as a Biological Variable (SABV) integration in the design, analysis, and reporting of clinical data (42,43). CIHR reports that they met their target of 67% of proposals addressing sex or gender considerations in the 2019-2020 grant cycle (44). CIHR applicants are required to answer yes or no (via a checkmark) to two questions on whether or not sex or gender is integrated into the proposal. An analysis of submitted funding to CIHR from 2011 to 2019 indicated that by 2019, 83% of submitted grants had checked yes to sex and 33% of submitted grants had checked yes to gender(33). It is likely that the earlier adoption of SGBA in Canada led to more reporting on the use of sex and gender considerations in research. However, it is also feasible that the mandatory nature of SGBA reporting boxes used in the analysis resulted in higher percentages of SGBA considerations at CIHR (39) relative to other funding agencies. In addition, given that the funding rates at CIHR are below 20%, it is also important to consider what percentage of funded rather than submitted proposals mention sex and gender in their abstracts. In the published literature, estimates of analyses of possible sex differences are lower at approximately 2-7% across a variety of disciplines (23,37,45) including papers in which the authors’ institutions were in Canada or the United States. Therefore, we sought to explore whether abstracts mentioned sex, gender, or studied historically underrepresented groups in health research (female, 2SLGBTQ+) within the publicly available abstracts of CIHR-funded operating and project grants.

In this paper, we analyzed publicly available data from the CIHR funding database from 2009-2020 to determine whether proposal abstracts across all areas of health research mentioned sex or gender in the abstract. To understand other health inequities, we also examined whether the abstract was focussed on research in female-specific health or 2S/LGBTQ+ health. We hypothesized that mentioning sex, gender, female or 2S/LGBTQ+ in the abstracts would increase over time as awareness improved and institutional requirements evolved. Further, we hypothesized that following CIHR’s SGBA mandates in 2019, 2020 abstracts would have increased mention of the population studied, including sex, gender, female specific or 2S/LGBTQ+ community members.

## Methods

We examined the published public abstracts of all Operating and Project Grants funded by the Canadian Institutes of Health Research (CIHR) from 2009 to 2020 using the CIHR Funding Decisions Database (46). These grants are akin to the R01 mechanism at NIH. From 2009-2015, CIHR held an Operating Grant program, which was renamed Project Grants in 2016. These programs are open competitions that fund investigator-driven research projects investigating fundamental and applied health research (47). In 2009, the average grant was just over $500,000 and in 2020, the average grant awarded was $751,022 over 4.39 years. To identify relevant grant abstracts, we used search terms “male, female, sex, gender, woman, women, uterus, uter, pregnan, breast, ovary, ovarian, ovaries, girl, boy, menopause, postpartum, maternal, placenta, prostate, cervi, testic, testes, vagin, penis, penial, binary, lesbian, gay, bisexual, MSM, FSF, queer, LGBT, LGBTQ, LGBTQIA, LGBTQIA+, LGB, transgender, transsex, trans, 2S, two spirit, 2 spirit, indigi (indigiqueer)” and then proceeded to manually code grant abstracts into appropriate categories (described below) based on the content of the abstract. For example, if an abstract mentioned sex, it was coded as sex mentioned. Additionally, grants proposing research across more than one category were double coded. For example, an abstract proposing investigating the effects of HIV during pregnancy on the mother and sex-specific effects on their children would be coded as both mentioning female-specific and sex differences. If the sex of the children was not mentioned, it would be coded as female-specific alone. Grant abstracts that did not clearly fall into one or more categories (n=12) were discussed between the three lead coding team members (ABN, TNS, TFLS) and were coded appropriately once a unanimous decision was met.

### Inclusion Criteria

All CIHR Operating (2009-2015) and Project (2016-2020) grant abstracts that were funded during the open competitions and awarded to Canadian institutions were included in our analysis (n= 8,964). Grant abstracts in both English (n=8,928) and French (n=36) were examined. French abstracts were translated using the Google translate tool, and then coded.

### Categorization of grant abstracts

Abstracts published with awarded grant information were examined and designated into the following categories (if relevant) based on the CIHR definition of SGBA: sex (any grant with an abstract that would mention sex: male, female, intersex), gender (any grant with an abstract that would mention gender (women, men, transgender, non-binary, gender non conforming, etc), female-specific grants (any grant with an abstract that would include only female participants or mention female-specific conditions), 2S/LGBTQ+ (any grant with an abstract that would include members of the 2S/LGBTQ+ population specifically or mention differences in outcomes for 2S/LGBTQ+ individuals compared to other populations). Any abstracts that did not fit in the above definitions were categorized as “sex/gender omitted”. Additionally, within the female-specific category, as breast cancer disproportionately receives more funding (15), we categorized female-specific cancer grant abstracts (any grant with an abstract that mentioned investigating cancer in a female-only population or investigating female-specific cancers) separately.

Given the limited methodological information contained in a grant abstract, we used a low threshold for categorization. To be included in our analysis, an abstract did not need to focus specifically on sex differences, gender differences, females, or 2S/LGBTQ+ individuals or explicitly state SGBA methodology; for example, mentioning the use of male and female rodents would have resulted in the abstract being coded into sex mentioned grant.

To understand whether the mention of sex, gender or female differed across research subjects we analyzed the CIHR public database across those funded abstracts or keywords that mentioned “human”, “mice” or cells (“cell culture”, “in vitro”, “cell lines”). We expressed the grant abstracts mentioning sex, gender or female as a function of total-funded abstracts using the categories of human, mice or cell.

### Statistical Analyses

For each category in any given year, percentages were calculated for both grant abstracts awarded and the amount of funding awarded as a function of the total number of grants awarded or total funding dollars disbursed within that year, respectively. To examine funding trends over time, we ran simple regression analyses on both the percentage of grant abstracts awarded and the percentage of funding dollars awarded across the twelve years (2009-2020). We present the data by binning the grant abstracts into 4 year intervals (2009-2012, 2013-2016, 2017-2020). As cancer research in female health (breast, ovarian, cervical, uterine, etc.) receives greater research funding than other female-specific health conditions (15,16), we did a sensitivity analysis by removing those grant abstracts from the overall female-specific analysis. Frequency of keyword use of sex or gender across humans, mice or cell studies were analyzed using a Chi-square analysis. Statistics were calculated using Statistica statistical analysis software (v. 9, StatSoft, Inc., Tulsa, OK, USA). Significance was set at α=0.05.

## Results

Overall, 8,964 Operating and Project Grants were awarded between 2009-2020, totaling $611,807,644 in research funding. Of these, 91.65% of grant abstracts were coded as “sex/gender omitted”. Of those remaining, the percentage of grant abstracts that included mention of: sex: 1.94%, gender:0.66%, female-specific: 5.92%, female-specific without cancer: 4.07%, and 2S/LGBTQ+: 0.35% (Figure 2a).

**Figure 2.**
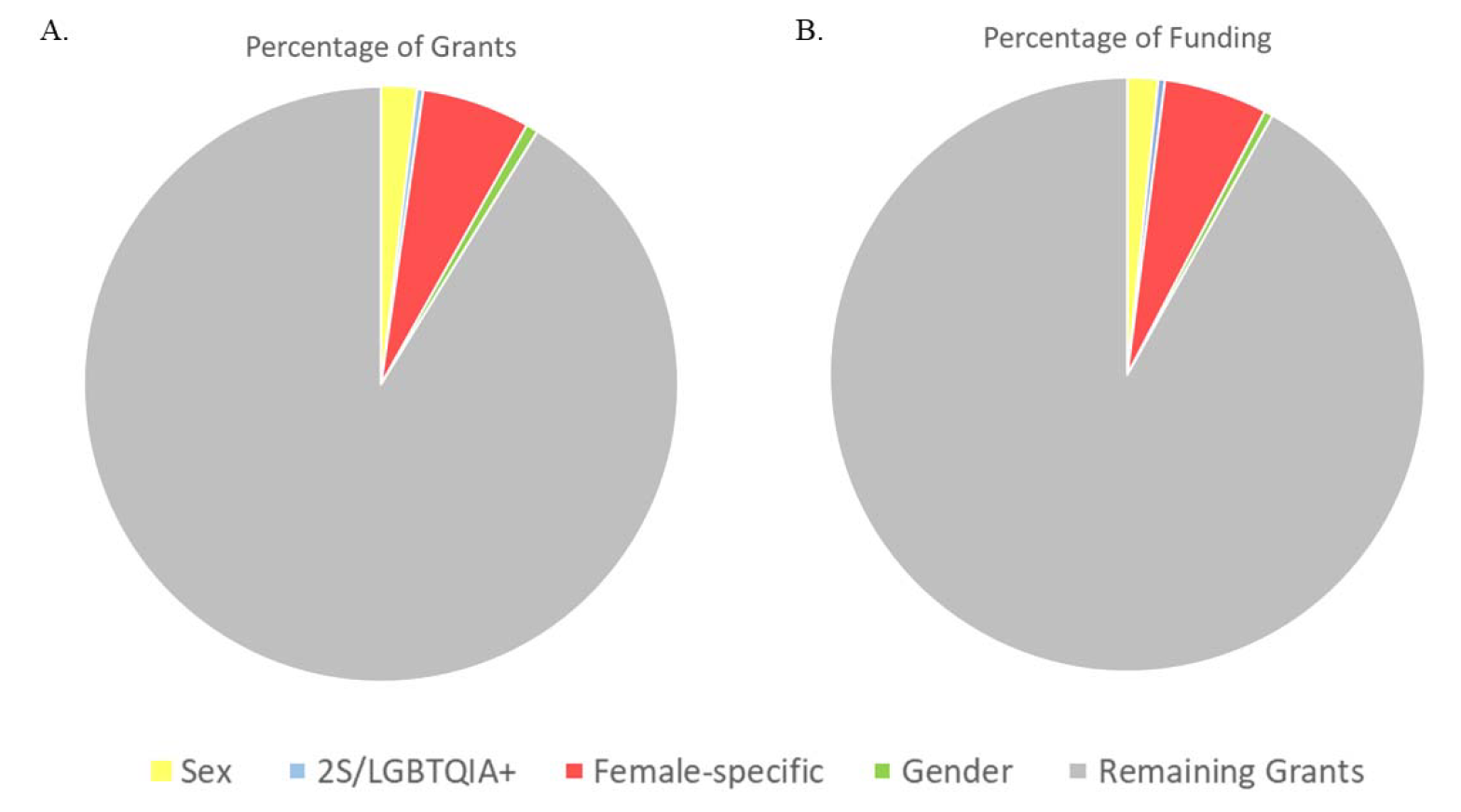
**Mean percentage of grant abstracts and funding awarded to each category of health research that mentioned the terms listed.** A) Pie chart of the percentages of Canadian Institutes of Health Research (CIHR) grants in each category (% of total grant abstracts) from 2009-2020. N=8,964. Sex : 1.94% (n=170); 2S/LGBTQ+: 0.35% (n=33); female-specific: 5.92% (n=517); gender : 0.66% (n=60); “Sex/Gender omitted”: 91.65% (n=8,230). Additionally, 4.07% of the grant abstracts were female-specific excluding cancer grants (n=367). B) Pie chart of the percentage of CIHR funding dollars (% of total funding amounts) for categorized grants abstracts from 2009- 2020. Sex: 1.67%; 2S/LGBTQ+: 0.36%; female-specific: 5.65%; gender: 0.46%; Sex/gender omitted: 92.27%. Additionally, 3.81% of the grants were female-specific excluding cancer grants.

### Both the percentage of grants with abstracts mentioning sex, and percentage of funding dollars for grant abstracts mentioning sex, more than doubled across 2009-2020

Overall, from 2009-2020, the percentage of funded grant abstracts that mentioned sex was 1.94% (Figure 2a). This percentage increased from 1.30% in 2009 to 2.86% in 2020 (R^2^ = 0.56, F (1, 10) = 12.91, p = 0.005, □ = 0.75). After the SGBA evaluation criteria were adopted by CIHR in 2019, grants with abstracts mentioning sex in the abstract in 2020 comprised 2.86% of funded grants, which was a decrease from 3.21% in 2018 and 4.43% in 2019.

Results indicated that grants with abstracts mentioning sex were awarded 1.67% of all funding dollars from 2009-2020 (Figure 2b), which increased significantly from 1.07% in 2009 to 2.33% in 2020 (R^2^ = 0.37, F (1, 10) = 5.99, p = 0.03, □ = 0.61).

### The percentage of grant with their abstracts mentioning gender did not change from 2009- 2020 and funding amounts decreased

Grants with abstracts mentioning gender were 0.66% of total CIHR-funded grants from 2009-2020 (Figure 2a) and 0.57% of projects funded in 2020. Although the percentage of grants that were funded decreased from 0.64% in 2009 to 0.57% in 2020 (see Figure 3), this was not statistically significant (R^2^ = 0.18, F (1, 10) = 2.20, p = 0.17, □ = -.042).

**Figure 3.**
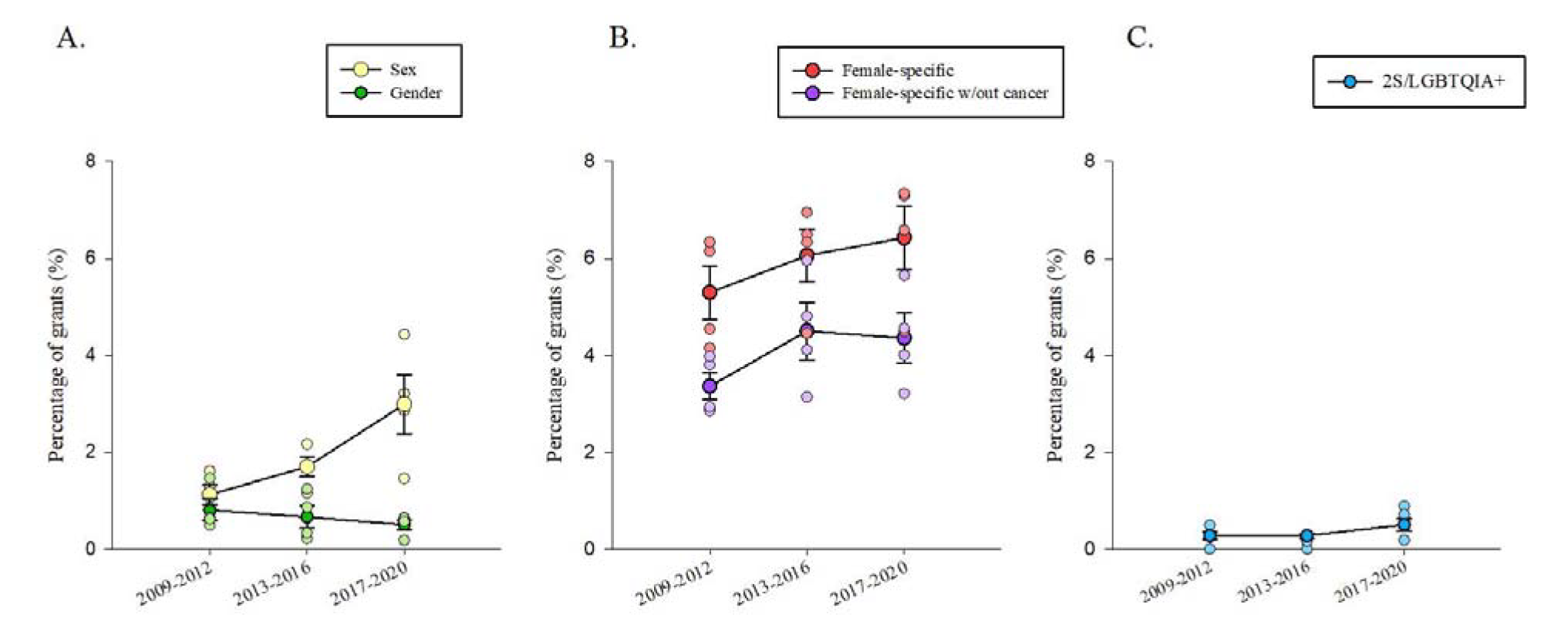
**Mean percentage of grants awarded to each category of health research.** A-C) Percentage of total grants funded by Canadian Institutes for Health Research (CIHR) binned in three time points: 2009-2012, 2013-2016, and 2017-2020. Data points indicate individual years within the range and error bars represent standard error of the mean (SEM). A) Percentage of grants funded by year that mentioned sex (yellow) and gender (green) in their abstracts remained below 2% throughout the years. Grant abstracts that mentioned sex increased over time (R^2^ = 0.56, F (1, 10) = 12.91, p = 0.005, □ = 0.75) but grant abstracts that mentioned gender did not (R^2^ = 0.18, F (1, 10) = 2.20, p = 0.17, □ = -.042). B) Percentage of grant funding amounts (funded by year) that mentioned female-specific health or disease factors (red), or with female-specific health without cancer research (purple). The percentage of grants awarded in either category did not increase over time (Female-specific: R^2^ = 0.08, F (1, 10) = 0.82, p = 0.39, □ = 0.28; Female-specific without cancer: R^2^ = 0.10, F (1, 10) = 1.15, p = 0.32, □ = 0.32). The percentage of grants that mentioned 2S/LGBTQ+ health (blue) in their abstracts funded by year remained below 1% across all years but increased significantly over time (R^2^ = 0.36, F (1, 10) = 5.53, p = 0.04, □ = 0.60).

**Figure 4:**
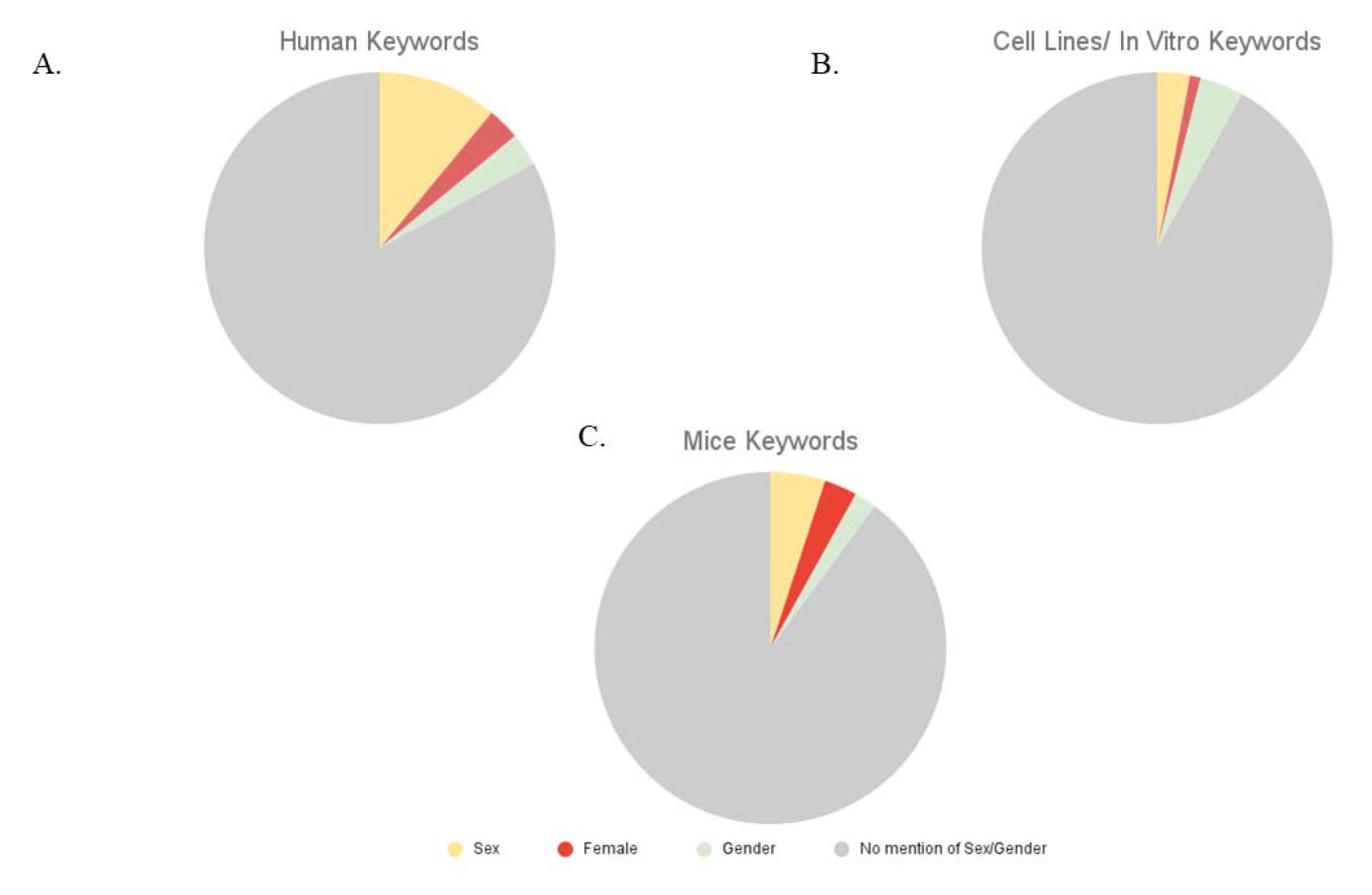
**Keyword search for mentions of sex, gender, or female health, in given proposal keywords separated by subject category.** Data obtained from search of “keywords” associated with grant abstract. (A) In grants with Human subjects (obtained by key word search): 11% (n=34) also had a keyword of “sex”, 3% (n=10) had a keyword of “female”, 3% (n=9) had a keyword of gender, and 83% had no keyword mention of sex, gender, or female health. (B) In grants with keywords that mentioned cell lines, cell culture or in vitro subjects: 3% (n=7) also had a keyword of “sex”, 1% (n=3) had a keyword of “female”, 4% (n=9) had a keyword of gender, and 92% had no keyword mention of sex, gender, or female health. (C) In grants with mice subjects (obtained by keyword search): 5% (n=37)) also had a keyword of “sex”, 3% (n=25) had a keyword of “female”, 2% (n=17) had a keyword of gender, and 89% had no keyword mention of sex, gender, or female health.

The percentage of funding dollars awarded to grants with abstracts mentioning gender remained below 1% (0.46%) of the total amount of funding awarded by CIHR from 2009- 2020 (Figure 2b). This percentage significantly decreased from 0.80% in 2009 to 0.31% in 2020 (R^2^ = 0.46, F (1, 10) = 8.55, p = 0.02, □ = −0.68). This indicates that although the percentage of grants awarded to study gender did not change, funding dollars awarded to such grants decreased over time (Table 1).

**Table 1.** Funding data in four-year bins. Total funding amounts in Canadian dollars and average percentage of total funding amount (%) for female-specific grants, female-specific grants without cancer grants (excluding any grant abstracts related to female-specific cancers), grants in which the abstracts mentioned sex, gender, or 2S/LGBTQ+ by year awarded by CIHR.

| **Years** | **Female-specific grants funding** | **Female-specific w/out cancer grants funding** | **Sex grants funding** | **Gender grants funding** | **2S/LGBTQ + grants funding** | **Total for all grants funding** |
| --- | --- | --- | --- | --- | --- | --- |
| **2009-2012** | \$88,176,847<br>(4.76%) | \$53,590,011<br>(2.91%) | \$19,471,838<br>(1.07%) | \$10,587,955<br>(0.58%) | \$4,689,111<br>(0.25%) | \$1,838,500,442 |
| **2013-2016** | \$97,552,417<br>(6.00%) | \$70,813,456<br>(4.55%) | \$21,339,244<br>(1.42%) | \$8,441,267<br>(0.50%) | \$4,782,071<br>(0.27%) | \$1,731,993,160 |
| **2017-2020** | \$123,992,008<br>(6.19%) | \$78,583,385<br>(3.97%) | \$53,585,969<br>(2.51%) | \$6,486,435<br>(0.31%) | \$12,219,729<br>(0.58%) | \$2,002,035,786 |
| **Total (2009-2020)** | \$309,721,272<br>(5.65%) | \$203,682,822<br>(3.81%) | \$94,397,051<br>(1.67%) | \$25,515,657<br>(0.46%) | \$21,690,911<br>(0.36%) | \$5,714,297,846 |

### Grant with abstracts mentioning female health did not change across 2009-2020, but the percentage of funding dollars increased by more than 3%

Our results indicate that from 2009-2020, 5.92% of funded grants were allocated to female-specific health research (Figure 2a). There was no significant difference in the percentage of grant abstracts that mentioned female-specific research over time (from 4.54% in 2009 to 6.58% in 2020) (R^2^ = 0.08, F (1, 10) = 0.82, p = 0.39, □ = 0.28). Furthermore, 5.65% of CIHR funding dollars were awarded to female-specific proposals (as identified by their abstracts) from 2009-2020 (Fig. 2b), which did significantly increase across the years from 3.37% in 2009 to 6.84% in 2020 (R^2^ = 0.32, F (1, 10) = 4.75, p = 0.05, □ = 0.57).

We next performed a sensitivity analysis by removing grants awarded for “female” cancers that were identified by their abstracts (15,16). After removing these grants, grants awarded to female-specific research comprised 4.07% of total grants awarded from 2009- 2020. There was no significant change in the percentage of female-specific grants, excluding cancer, over time (from 2.85% in 2009 to 4.01% in 2020) (R^2^ = 0.10, F (1, 10) = 1.15, p = 0.32, □ = 0.32; Figure 3b).

However, removing cancer grants did impact funding trends over time. The percentage of funding allocated to female-specific health, excluding cancer research, fell to 3.81% from 2009-2020 (Table 1), and the difference in funding dollars awarded across years for female-specific health was no longer significant (1.95% in 2009 to 4.00% in 2020) (R^2^ = 0.12, F (1, 10) = 1.34, p = 0.27, □ = 0.34). This suggests that research on female-specific cancers was driving the increase in funding for female-specific health between 2009 and 2020.

### The percentage of grant abstracts mentioning 2S/LGBTQ+-specific health and funding allocated to these projects more than tripled across 2009-2020, but remained less than 1% of all funded grants

Grant abstracts that mentioned 2S/LGBTQ+-specific health comprised less than 1% (0.35%) of total grants funded by CIHR from 2009-2020 (Figure 2a). There was a significant increase in the percentage of grants funded throughout the years, from 0.13% in 2009 to 0.43% in 2020 (R^2^ = 0.36, F (1, 10) = 5.53, p = 0.04, □ = 0.60; Figure 3c). The percentage of grant money awarded to grants mentioning 2S/LGBTQ+-specific health also remained below 1% (0.36%) from 2009-2020 (Figure 2b), increasing from 0.05% in 2009 to 0.46% in 2020, which was not significant (R^2^ = 0.22, F (1, 10) = 2.82, p = 0.12, □ = 0.47).

### The percentage of grants funded and funding dollars awarded to grant that omitted mention of sex/gender in their abstracts decreased by roughly 3% across 2009-2020

Grants that had abstracts that did not mention sex/gender (did not mention sex, gender, female specific or 2S/LGBTQ+ specific health) comprised 91.65% of total grants overall from 2009-2020 (Figure 2a). This percentage significantly decreased over time, from 93.51% in 2009 to 90.41% in 2020 (R^2^ = 0.41, F (1, 10) = 7.07, p = 0.02, □ = −0.64; data not shown). Grants with abstracts that did not mention sex/gender received 92.27% of funding dollars from 2009-2020 (Figure 2b). This percentage also decreased significantly over time, from 93.58% in 2009 to 91.09% in 2020 (R^2^ = 0.46, F (1, 10) = 8.49, p = 0.02, □ = −0.68).

### No significant differences seen in the proportion of funded grant keywords using sex, gender, or female across “subject type”

Subject population is not a searchable field on the CIHR database. Thus we searched abstracts and keyword fields for the search terms listed in the Methods section. We performed a chi-square analysis to determine whether the percentage of funded grants using abstracts and keywords that mentioned the populations to be studied (human (n=310), mice (n=748), cells(n=236)) and cross-referenced with mention of sex, gender or female. We combined the terms sex or gender for mice and cells in the analysis due to conflation of these terms in animal models. The percentage of funded projects with sex/gender in keywords was not significantly different across population groups (χ^2^= 4.784, p=0.091).

## Discussion

Overall, our analysis of CIHR research funding allocation by reviewing the awarded abstracts revealed that, from 2009-2020, 91.65% of grants did not have sex or gender mentioned in their abstract of proposed research. Across all groups examined, the percentage of grants awarded and the funding dollars allocated to projects which in their abstracts mentioned female-specific health, 2S/LGBTQ+-specific health, or sex or gender of the research participants was below 10%, and slightly lower for funding dollars compared to percentage of grants funded. Although the percentage of grants with abstracts that mentioned sex or 2S/LGBTQ+-specific health doubled over the 12 years, both remained under 3% of grants funded in 2020 (Figure 5a). The percentage of CIHR grants with abstracts that mentioned female-specific health or gender in health did not significantly change across the years. However, the percentage of grants with abstracts that did not mention sex/gender declined by approximately 3% over the twelve years yet still remained at more than 90% of grants in 2020. Overall, we found that CIHR mandatory SGBA reporting in applications in 2010 and scoring in 2019 was commensurate with an increase in the amount of funded health research that mentioned two of the specified groups in our analysis (2S/LGBTQ+ and sex; see Figure 5b). Although the gains have been modest (increasing by 0.3% and 1.5%, respectively), our findings suggest that overall funding allocated to grants mentioning sex, gender, female-specific health, or 2S/LGBTQ+-specific health research remains below 10% of all funded Project/Operating grants at CIHR after more than 10 years of SGBA adoption.

**Figure 5.**
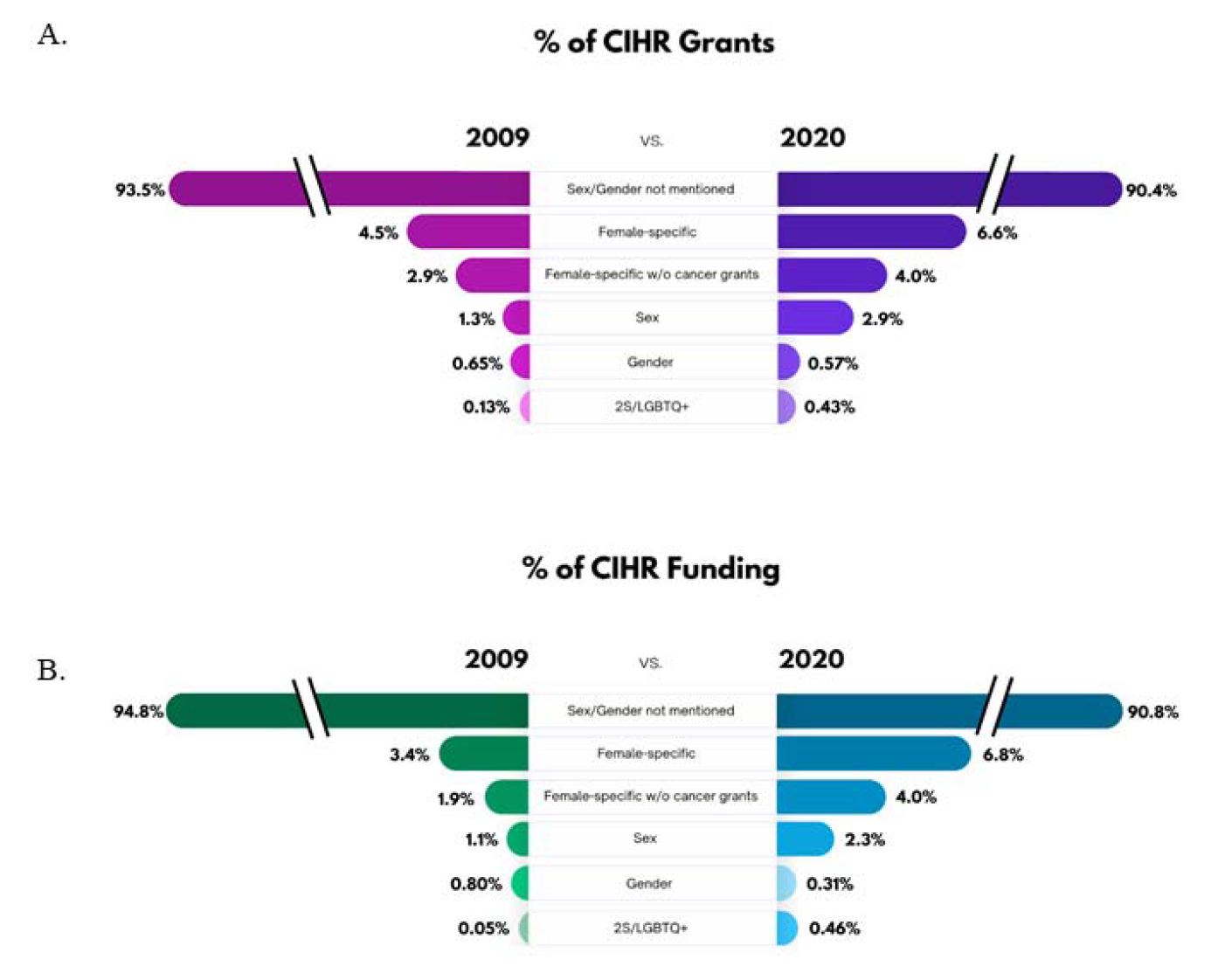
**An infographic depicting the change in percentage of grants and funding between 2009 and 2020 for awarded Canadian institutes of Health Research (CIHR) grants for the different categories.** The change in percentage (%) of grants (A) and funding amount (B) in the years 2009 and 2020 that did not mention of sex and gender in their grant abstracts or mentioned female-specific health, female-specific health not including cancer based grants, sex, gender, or 2S/LGBTQ+ health.

### The percentage of funded grants with abstracts mentioning sex or gender has largely remained unchanged and funding amounts have stagnated or declined

In the current study, we found that funded projects mentioning sex or gender in their abstracts comprised 2.57% of grant abstracts funded and 2.13% of the research dollars dispersed from 2009-2020. Although the percentage of grants with abstracts mentioning sex increased significantly by 2% between 2009 and 2020, the percentage of grants with mentions of gender in the abstract did not change across the years. Moreover, when comparing the percentage of grants with abstracts mentioning sex with the amount of funding allocated to these projects, we found that funding amounts only increased by 50% and was disproportionately smaller (2.68% versus 3.12% of grants), indicating funding amounts are increasing at a slower rate than the percentage of grants awarded. In contrast, funding for grants that mentioned gender in their abstracts decreased across this same time period.

Worldwide, research funding bodies have recognized the importance of bridging the gap in knowledge of sex and gender differences in health and implemented mandates to encourage its scientific inquiry. An analysis of CIHR submitted grants reported that in 2019, 83% of submitted grants had checked that they were considering sex and 33% considered gender in their research (33). However, our results examining the published funded abstracts, suggest a different conclusion, albeit with several caveats due to the nature of the data we can publicly assess. The differences in findings between the current analyses and those from Haverfield and Tannenbaum (2021) may be due the population of grants analyzed (funded versus submitted, respectively) or in what was analyzed (public abstract versus mandatory checkbox). It will be important to understand whether or not future publications stemming from these funded projects result in dissemination of SGBA. However, the current findings of a low percentage of mention of sex/gender in the abstracts are in line with previous reports that demonstrate that publications in a wide variety of disciplines have low rates of dissemination and analyses of findings by sex or gender (23,24,48,49). It is very possible that researchers did not include sex/gender in their abstracts but will ultimately publish their findings with respect to sex/gender. To understand the relationship between funding and eventual publications, it would be important to have a public repository of published findings linked to the funded projects, similar to that of other funding agencies, such as NIH.

### Cancer research received more money per grant than other female-specific health grants

In the current study, we found that female-specific research comprised 5.92% of grants funded from 2009-2020 and correspondingly 5.65% of research dollars disbursed, and neither measure increased significantly over time. Female-specific cancer research accounted for approximately 31.3% of this funding; removing these grants we found that 4.07% of grants that were funded mentioned female-specific health in their abstract. This draws a parallel to previous findings in the published literature, where only 5% of neuroscience publications (50), 3-4% of neuroscience and psychiatry papers (23) or under 1% of human imaging studies investigated female-specific health (51). This is disproportionately lower than male-specific studies that represented up to 50% of neuroscience publications in 2010- 2014 (50), and 27% of neuroscience and psychiatry publications in 2009 and 2019 (23). These findings are important as data from the United States shows that diseases that disproportionately affect women are less likely to be funded (14). Taken together, these findings suggest other efforts will need to be adopted to improve women’s health and female-specific health funding.

Gynecologic and breast cancer grants (as identified by abstracts) accounted for approximately 31% of the female-specific funded grants. This suggests that female cancer research initiatives received a third of the overall amount of funding dollars awarded for female-specific health inquiries. To make progress towards health equity, it is important for funders and researchers to recognize that many diseases have unique risks, symptomatology and treatment in females and that female-unique experiences alter health outcomes and disease risk (11,25,52,53). For example, disorders during pregnancy, such as preeclampsia and gestational hypertension elevate the risk of type 2 diabetes and cardiovascular disease later in life (52). Despite evidence of female-specific factors influencing health conditions (54), studies have neglected the adequate use of females in their research (25), contributing to poor health outcomes in females, including greater adverse side effects to new therapeutic drugs (11,53).

### 2S/LGBTQ+-Specific Health was awarded less than 1% of the funding dollars

Our findings indicate that grants with abstracts mentioning 2S/LGBTQ+-specific health comprised less than 1% of overall funded grants and funding dollars from 2009-2020 at CIHR for Operating and Project Grants. Although the proportion of grants with abstracts mentioning 2S/LGBTQ+-specific health did increase across the 12 years (0.3%), the amount of funding dollars did not significantly increase across the same time. 2S/LGBTQ+-specific health has been systematically excluded from health research resulting in greater health disparities for 2S/LGBTQ+ community members (55). For example, lesbian women are less likely to access cancer prevention services compared to heterosexual women (56) and gay men are at a higher risk of contracting HIV compared to their heterosexual counterparts, especially among communities of color (34). Furthermore, 2S/LGBTQ+ individuals are at higher risk for poor mental health outcomes (57), psychological distress (58), and suicidal ideation (59) relative to heterosexuals. 2S/LGBTQ+ community members have higher rates of disability (35) and poorer general health (59). Thus, 2S/LGBTQ+ health requires explicit attention and funding dollars to address these disparities. Although it is encouraging that 2S/LGBTQ+ research is trending in the right direction, dedicated funding initiatives are required to help close the health disparity gap for the 2S/LGBTQ+ community.

### Limitations

Although the current study contributes to our understanding of the funding landscape with respect to successful grants mentioning sex, gender, women and 2S/LGBTQ+ health in Canada, there are limitations to this study. First, it is important to consider we were only able to look at publicly available abstracts (which is limited to a maximum of 2000 characters) rather than the full scientific summaries or the grant proposals themselves. The character constraint of these public abstracts introduces a number of limitations, and having access to full summaries may have allowed us to evaluate SGBA intentions in CIHR Operating and Project grant proposals outside of what was examined in this study, mention of sex or gender in the study population.

As we only examined the public abstracts of grants, in the future it would be important to examine the concordance of the public abstract to the scientific abstracts and/or the full grant proposals themselves. It is very possible that grant proposals were using sex and gender but did not record the use of their population in the abstract. It also seems reasonable to expect that if sex and gender considerations were central to the experimental design, descriptions of the participants by sex and gender would have been mentioned in the abstract. However, it is possible this assumption is not correct. An informal analysis by the authors suggest a great variability in concordance of the abstract to the grant proposal itself with respect to sex and gender as LAMG found 100% concordance of mention of sex/gender in the abstract to submitted grants as principal investigator, 90% as co-investigator but only 33% as a reviewer. In the future it is our recommendation that public funders, such as CIHR, ensure that there is publicly available data on the population studied within the grant proposal (human, animal, cell lines) and whether or not sex/gender, female health or 2SLGBTQ+ health will be considered.

Another important point for funding agencies to consider is what amount of information in the proposal itself would constitute compliance with the SGBA mandate. Is a one sentence mention of an analysis by sex and/or gender in the grant enough? Should the hypotheses, background information and methods sections consider sex and/or gender? Clarifying how to achieve the goal of the SGBA mandate would enable researchers to better comply with the mandate and allow for more in-depth analyses of compliance to be completed. We also recommend that CIHR include the type of subject used per proposal (ie, human, mice, cell lines etc), to be included in the publicly available database.

Results in the present paper mirror those of analyses examining the published literature (23,37,45,50). The present study found that under 4% of proposal abstracts mention sex or gender and ∼5% mention female-specific researchers which aligns with reports in the published literature across multiple disciplines (23,60). In terms of studies statistically examining a possible sex difference in neuroscience or psychiatry, one study found approximately 5% of neuroscience studies examined sex as a possible discovery variable (rodent, human, cells studies were considered; (23)) and another study found 14% of human psychiatric studies (61). Across a wide variety of disciplines (behavior, behavioral physiology, endocrinology, general biology, immunology, neuroscience, pharmacology, physiology and reproduction), approximately 7% of studies considered sex as a discovery variable (60). All of the above mentioned studies excluded the use of sex as a covariate as covarying sex is problematic on a number of levels (62). When removing studies that used sex as a covariate or other non-optimal analyses (not including sex as a discovery variable), the percentage of publications using sex in the analyses was 2-7% of publications across a variety of disciplines (23,60), which is at least partially consistent with our results. We also see alignment with our finding of approximately 5% of funded grant abstracts referring to female-specific research and publications finding 0.5-5% of a wide variety of research is in females (23,51). However, it is also important to note that microarray and RNA-seq databases have more equivalent use across males and females (females: 21.6%-25.8% (human and mice) versus males: 18.9%-31% (63).

Further, although we examined nearly 9000 Canadian CIHR project grants, it is crucial to understand worldwide health research funding trends. Future research should consider evaluating other funding agencies’ mention of sex/gender in project abstracts, such as the NIH, Horizon Europe, the Council of Scientific and Industrial Research (CSIR, India), Nottingham China Health Institute (NCHI), and the African Academy of Sciences. To this end, other analyses of disease burden would also be important to uncover, as one study found that funding was disproportionately lower for diseases that had a greater burden for females (14).

### Perspectives and Significance

It is important to underscore that much of the existing work in this field has been focused on binary sex and gender. Looking beyond this allows for a more nuanced understanding of complex issues that include multiple perspectives. Given the importance of studying sex, gender, females-specific factors, and 2S/LGBTQ+-specific effects on all four pillars of research (biomedical, clinical, population health, and health services), we have developed recommendations for future promotion and evaluation of sex and gender in health research. First, we recommend specific, ring-fenced funding for sex, gender, female-specific health, and 2S/LGBTQ+-specific health. Dedicated funding has a multifold effect to increase the number of investigators, and the number of publications which drives discovery into disease management and treatments. Recent examples of this can be seen with research funding awarded to AIDS and ALS (64,65). Indeed, CIHR, via funding initiated directly from the Institute of Gender and Health, has held successful funding competitions dedicated to Sex as a Biological variable. In 2017, this competition awarded just over $4.3 million in Catalyst Grants for Sex as a Variable in Biomedical Research. However, this represents 3.36% of funding for specific research areas awarded by Institute-driven research competitions (e.g. catalyst and other grants) in that same year. Although the success of this funding competition was modest in the context of CIHR’s larger Project Grant budget, it is possible these Catalyst funds spurred the increase in grants on sex differences in 2018-2020 as seen in Figure 3A. Increasing dedicated funding amounts and requests for applications to reflect the need for health equity, greater engagement in sex, gender, female-specific health, and 2S/LGBTQ+- specific health would not only encourage future research proposals to properly integrate SGBA, but also inspire the next generation of researchers to investigate the role of sex and gender in health, female-specific health, and 2S/LGBTQ+ specific health.

For many granting agencies, once funds are awarded there is no reporting mechanism in place to determine if the awarded money was spent on the proposed project. Therefore, despite requiring a disclosure of how SGBA is represented in the grant application, there is no requirement for performing research with appropriate sex or gender considerations to receive funding, nor is there a requirement to report on SGBA findings at the end of the award. Funding agencies should consider introducing mechanisms of accountability for SGBA after funding has been awarded (48).

Publishers also play a critical role in advancing sex and gender in health research. An academic publisher is tasked with the proper validation of scientific findings (66). As such, implementing rigorous standards for SGBA is an important step to enhance the uptake and utilization of SGBA. For example, as of June 2022, researchers who submit to certain Nature Journals are prompted to state why or why not SGBA was used in addition to noting if results are gender-or sex-specific in the title or abstract (67). Furthermore, the Sex and Gender Equity in Research (SAGER) guidelines have been developed to ensure authors supply information on sex and gender throughout their manuscripts, including in the title and abstract (68). These additions monitor SGBA at an additional step in the academic research and dissemination process. Funding agencies could also adopt a similar approach, to require the grant abstracts include population characteristics with respect to sex and gender to make it more salient to the public what population is being examined.

Although collectively the understanding of the need for and recognition of sex and gender in research may be improving, it still remains at low levels in publications and dissemination and perhaps funding. The research community needs to acknowledge that for publications, clinical trials, and grant funding applications, SGBA integration needs improvement. In line with previous reports (23,29,67), we found that grants awarded to research mentioning sex and gender differences, female-specific health, and 2S/LGBTQ+- specific health consistently represented a small percentage of awarded grants and funding from 2009-2020. The aspiration of SGBA is that widespread adoption of its principles will result in an equitable future for health where female-specific health, 2S/LGBTQ+ health, racialized and gendered experiences, and more are considered. SGBA mandates and frameworks are a first step in ensuring researchers contribute to a more representative body of knowledge, but continued efforts are needed to improve knowledge dissemination and increase funding to advance health equity in research.

Sex and gender meaningfully contribute to differences in health through differing disease risk and manifestation, treatment response, and healthcare seeking behavior (1,10,26,27). However, sex and gender have been historically neglected in health research, leaving gaps in our knowledge of how best to diagnose and treat illnesses affecting people of both sexes and any gender identity. These gaps contribute to ongoing sex and gender disparities in health outcomes that may begin to be rectified if more time and resources are devoted to such research.

## List of Abbreviations

SGBA: Sex and Gender-based Analysis
CIHR: Canadian Institutes of Health Research
NIH: National Institute of Health
SABV: Sex as a Biological Variable
2S/LGBTQ+: Acronym used to signify Two-Spirit, Lesbian, Gay, Bisexual, Transgender, Queer or Questioning, Intersex, and Asexual people collectively.

## Declarations

### Ethics approval and consent to participate

Not applicable

### Consent for publication

Not applicable

### Availability of data and material

All material was from publicly available databases but is available upon request.

### Authors’ contributions

TNS, TFLS, ABN data collection and first draft; LAMG, KNM, TNS, ABN, TFLS study design, methodology, and editing.

## Acknowledgements

The authors thank Kenzie Hoffman for designing the Figure 5 infographic.

## References

1. Mauvais-Jarvis F, Bairey Merz N, Barnes PJ, Brinton RD, Carrero JJ, DeMeo DL, et al. Sex and gender: modifiers of health, disease, and medicine. Lancet Lond Engl. 2020;396(10250):565–82.

2. Rich-Edwards JW, Kaiser UB, Chen GL, Manson JE, Goldstein JM. Sex and Gender Differences Research Design for Basic, Clinical, and Population Studies: Essentials for Investigators. Endocr Rev. 2018 Aug 1;39(4):424–39.

3. Seney ML, Huo Z, Cahill K, French L, Puralewski R, Zhang J, et al. Opposite Molecular Signatures of Depression in Men and Women. Biol Psychiatry. 2018 Jul 1;84(1):18–27.

4. Mersha TB, Martin LJ, Biagini Myers JM, Kovacic MB, He H, Lindsey M, et al. Genomic architecture of asthma differs by sex. Genomics. 2015 Jul;106(1):15–22.

5. Barbieri D, García Cazorla A, Thil L, Mollard B, Ochmann J, Peciukonis V, et al. EIGE- 2021 Gender Equality Index 2021 Report. 2021;

6. Kessler RC, Bromet EJ. The epidemiology of depression across cultures. Annu Rev Public Health. 2013;34:119–38.

7. Loke H, Harley V, Lee J. Biological factors underlying sex differences in neurological disorders. Int J Biochem Cell Biol. 2015 Aug;65:139–50.

8. Häfner H, Riecher-Rössler A, Maurer K, Fätkenheuer B, Löffler W. First onset and early symptomatology of schizophrenia. A chapter of epidemiological and neurobiological research into age and sex differences. Eur Arch Psychiatry Clin Neurosci. 1992;242(2– 3):109–18.

9. Wada H, Miyauchi K, Daida H. Gender differences in the clinical features and outcomes of patients with coronary artery disease. Expert Rev Cardiovasc Ther. 2019 Feb;17(2):127–33.

10. Westergaard D, Moseley P, Sørup FKH, Baldi P, Brunak S. Population-wide analysis of differences in disease progression patterns in men and women. Nat Commun. 2019 Dec;10(1):666.

11. Zucker I, Prendergast BJ. Sex differences in pharmacokinetics predict adverse drug reactions in women. Biol Sex Differ. 2020 Jun 5;11(1):32.

12. Woodward M. Cardiovascular Disease and the Female Disadvantage. Int J Environ Res Public Health. 2019 Apr 1;16(7):E1165.

13. Mozaffarian D, Benjamin EJ, Go AS, Arnett DK, Blaha MJ, Cushman M, et al. Heart disease and stroke statistics--2015 update: a report from the American Heart Association. Circulation. 2015 Jan 27;131(4):e29-322.

14. Mirin AA. Gender Disparity in the Funding of Diseases by the U.S. National Institutes of Health. J Womens Health. 2021 Jul;30(7):956–63.

15. Coronado AC, Finley C, Badovinac K, Han J, Niu J, Rahal R. Discrepancies between Canadian cancer research funding and site-specific cancer burden: a spotlight on ten disease sites. Curr Oncol. 2018 Oct;25(5):338–41.

16. Carter AJ, Nguyen CN. A comparison of cancer burden and research spending reveals discrepancies in the distribution of research funding. BMC Public Health. 2012 Jul 17;12:526.

17. Galea LA, Qiu W, Duarte-Guterman P. Beyond sex differences: short and long-term implications of motherhood on women’s health. Curr Opin Physiol. 2018 Dec 1;6:82–8.

18. Rytz CL, Kochaksaraei GS, Skeith L, Ronksley PE, Dumanski SM, Robert M, et al. Menstrual Abnormalities and Reproductive Lifespan in Females with CKD: A Systematic Review and Meta-Analysis. Clin J Am Soc Nephrol [Internet]. 2022 Nov 23 [cited 2022 Nov 28]; Available from: https://cjasn.asnjournals.org/content/early/2022/11/20/CJN.07100622

19. Garcia M, Miller VM, Gulati M, Hayes SN, Manson JE, Wenger NK, et al. Focused Cardiovascular Care for Women: The Need and Role in Clinical Practice. Mayo Clin Proc. 2016 Feb;91(2):226–40.

20. Silveyra P, Fuentes N, Rodriguez Bauza DE. Sex and Gender Differences in Lung Disease. Adv Exp Med Biol. 2021;1304:227–58.

21. Golden LC, Voskuhl R. The importance of studying sex differences in disease: The example of multiple sclerosis. J Neurosci Res. 2017 Jan 2;95(1–2):633–43.

22. Yakerson A. Women in clinical trials: a review of policy development and health equity in the Canadian context. Int J Equity Health. 2019 Apr 15;18(1):56.

23. Rechlin RK, Splinter TFL, Hodges TE, Albert AY, Galea LAM. An analysis of neuroscience and psychiatry papers published from 2009 and 2019 outlines opportunities for increasing discovery of sex differences. Nat Commun. 2022 Dec;13(1):2137.

24. Geller SE, Koch AR, Roesch P, Filut A, Hallgren E, Carnes M. The More Things Change, the More They Stay the Same: A Study to Evaluate Compliance With Inclusion and Assessment of Women and Minorities in Randomized Controlled Trials. Acad Med J Assoc Am Med Coll. 2018 Apr;93(4):630–5.

25. Liu KA, Mager NAD. Women’s involvement in clinical trials: historical perspective and future implications. Pharm Pract. 2016;14(1):708.

26. Regitz-Zagrosek V. Sex and gender differences in health. EMBO Rep. 2012 Jul;13(7):596–603.

27. Thompson AE, Anisimowicz Y, Miedema B, Hogg W, Wodchis WP, Aubrey-Bassler K. The influence of gender and other patient characteristics on health care-seeking behaviour: a QUALICOPC study. BMC Fam Pract. 2016 Mar 31;17:38.

28. Ruben MA, Livingston NA, Berke DS, Matza AR, Shipherd JC. Lesbian, Gay, Bisexual, and Transgender Veterans’ Experiences of Discrimination in Health Care and Their Relation to Health Outcomes: A Pilot Study Examining the Moderating Role of Provider Communication. Health Equity. 2019 Jul;3(1):480–8.

29. Institute of Medicine (US) Committee on Lesbian G. The Health of Lesbian, Gay, Bisexual, and Transgender People [Internet]. National Academies Press (US); 2011 [cited 2022 Aug 16]. Available from: https://www.ncbi.nlm.nih.gov/books/NBK64806/

30. Reisner SL, Bradford J, Hopwood R, Gonzalez A, Makadon H, Todisco D, et al. Comprehensive Transgender Healthcare: The Gender Affirming Clinical and Public Health Model of Fenway Health. J Urban Health. 2015 Jun 1;92(3):584–92.

31. Keuroghlian AS, Ard KL, Makadon HJ, Keuroghlian AS, Ard KL, Makadon HJ. Advancing health equity for lesbian, gay, bisexual and transgender (LGBT) people through sexual health education and LGBT-affirming health care environments. Sex Health. 2017 Feb 6;14(1):119–22.

32. Clark BA, Veale JF, Townsend M, Frohard-Dourlent H, Saewyc E. Non-binary youth: Access to gender-affirming primary health care. Int J Transgenderism. 2018 Apr 3;19(2):158–69.

33. Haverfield J, Tannenbaum C. A 10-year longitudinal evaluation of science policy interventions to promote sex and gender in health research. Health Res Policy Syst. 2021 Jun 15;19(1):94.

34. Wolitski, Richard J., Ron Stall, and Ronald O. Valdiserri,. Unequal opportunity: health disparities affecting gay and bisexual men in the United States. USA: Oxford University Press; 2008.

35. Fredriksen-Goldsen, Karen I., Hyun-Jun Kim, and Susan E. Barkan. Disability among lesbian, gay, and bisexual adults: Disparities in prevalence and risk. Am J Public Health. 2012;102(1):16–21.

36. Woitowich NC, Woodruff TK. Implementation of the NIH Sex-Inclusion Policy: Attitudes and Opinions of Study Section Members. J Womens Health 2002. 2019 Jan;28(1):9–16.

37. Mamlouk GM, Dorris DM, Barrett LR, Meitzen J. Sex bias and omission in neuroscience research is influenced by research model and journal, but not reported NIH funding. Front Neuroendocrinol. 2020 Apr;57:100835.

38. McCarthy L, Milne E, Waite N, Cooke M, Cook K, Chang F, et al. Sex and gender-based analysis in pharmacy practice research: A scoping review. Res Soc Adm Pharm. 2017 Nov 1;13(6):1045–54.

39. Brady E, Nielsen MW, Andersen JP, Oertelt-Prigione S. Lack of consideration of sex and gender in COVID-19 clinical studies. Nat Commun. 2021 Dec;12(1):4015.

40. Zeraatkar D, Pitre T, Leung G, Cusano E, Agarwal A, Khalid F, et al. Consistency of covid-19 trial preprints with published reports and impact for decision making: retrospective review. BMJ Med [Internet]. 2022 Oct 1 [cited 2022 Oct 6];1(1). Available from: https://bmjmedicine.bmj.com/content/1/1/e000309

41. Directorate-General for Research and Innovation (European Commission). In-Depth Interim Evaluation of Horizon 2020. European Union; 2018 Oct.

42. Arnegard ME, Whitten LA, Hunter C, Clayton JA. Sex as a Biological Variable: A 5-Year Progress Report and Call to Action. J Womens Health. 2020 Jun;29(6):858–64.

43. Temkin SM, Barr E, Moore H, Caviston JP, Regensteiner JG, Clayton JA. Chronic conditions in women: the development of a National Institutes of health framework. BMC Womens Health. 2023 Apr 6;23(1):162.

44. Government of Canada CI of HR. 2022-2023 Departmental Plan [Internet]. Report No.: 2371–6827. Available from: https://cihr-irsc.gc.ca/e/52738.html

45. Woitowich NC, Beery A, Woodruff T. A 10-year follow-up study of sex inclusion in the biological sciences. Sugimoto C, Rodgers P, Shansky R, Schiebinger L, editors. eLife. 2020 Jun 9;9:e56344.

46. Government of Canada CI of HR. Funding Decisions Database [Internet]. 2008 [cited 2022 Aug 4]. Available from: https://webapps.cihr-irsc.gc.ca/decisions/p/main.html?lang=en#sort=namesort%20asc&start=0&rows=20

47. Government of Canada CI of HR. Project Grant Program [Internet]. 2022 [cited 2022 Oct 31]. Available from: https://cihr-irsc.gc.ca/e/49051.html

48. Shansky RM, Murphy AZ. Considering sex as a biological variable will require a global shift in science culture. Nat Neurosci. 2021 Apr;24(4):457–64.

49. Galea LAM, Choleris E, Albert AYK, McCarthy MM, Sohrabji F. The promises and pitfalls of sex difference research. Front Neuroendocrinol. 2020 Jan;56:100817.

50. Will TR, Proaño SB, Thomas AM, Kunz LM, Thompson KC, Ginnari LA, et al. Problems and Progress regarding Sex Bias and Omission in Neuroscience Research. eNeuro. 2017 Nov 9;4(6):ENEURO.0278-17.2017.

51. Taylor CM, Pritschet L, Jacobs EG. The scientific body of knowledge – Whose body does it serve? A spotlight on oral contraceptives and women’s health factors in neuroimaging. Front Neuroendocrinol. 2021 Jan 1;60:100874.

52. Zhao G, Bhatia D, Jung F, Lipscombe L. Risk of type 2 diabetes mellitus in women with prior hypertensive disorders of pregnancy: a systematic review and meta-analysis. Diabetologia. 2021 Mar;64(3):491–503.

53. Hurwitz N. Predisposing Factors in Adverse Reactions to Drugs. Br Med J. 1969 Mar 1;1(5643):536–9.

54. Galea LAM, Lee BH, de leon RG, Rajah MN, Einstein G. Chapter 45 - Beyond sex and gender differences: The case for women’s health research. In: Legato MJ, editor. Principles of Gender-Specific Medicine (Fourth Edition) [Internet]. Academic Press; 2023 [cited 2023 May 16]. p. 699–711. Available from: https://www.sciencedirect.com/science/article/pii/B9780323885348000456

55. Gorczynski PF, Brittain DR. Call to Action: The Need for an LGBT-Focused Physical Activity Research Strategy. Am J Prev Med. 2016 Oct;51(4):527–30.

56. Diamant, A. L., Schuster, M. A., & Lever, J. Receipt of preventive health care services by lesbians. Am J Prev Med. 19(3):141–8.

57. Dilley, J. A., Simmons, K. W., Boysun, M. J., Pizacani, B. A., & Stark, M. J. Demonstrating the importance and feasibility of including sexual orientation in public health surveys: health disparities in the Pacific Northwest. Am J Public Health. 100(3):460–7.

58. Chae, David H., and George Ayala. Sexual orientation and sexual behavior among Latino and Asian Americans: Implications for unfair treatment and psychological distress. J Sex Res. 2010;47(5):451–91.

59. Conron, Kerith J., Matthew J. Mimiaga, and Stewart J. Landers. A population-based study of sexual orientation identity and gender differences in adult health. Am J Public Health. 2010;100(10):1953–60.

60. Garcia-Sifuentes Y, Maney DL. Reporting and misreporting of sex differences in the biological sciences. Allison DB, Zaidi M, Vorland CJ, Kahathuduwa C, editors. eLife. 2021 Nov 2;10:e70817.

61. Duffy KA, Epperson CN. Evaluating the evidence for sex differences: a scoping review of human neuroimaging in psychopharmacology research. Neuropsychopharmacology. 2022 Jan;47(2):430–43.

62. Miller GA, Chapman JP. Misunderstanding analysis of covariance. J Abnorm Psychol. 2001 Feb;110(1):40–8.

63. Flynn E, Chang A, Altman RB. Large-scale labeling and assessment of sex bias in publicly available expression data. BMC Bioinformatics. 2021 Mar 30;22(1):168.

64. Ice Bucket Challenge dramatically accelerated the fight against ALS [Internet]. The ALS Association. [cited 2022 Oct 12]. Available from: https://www.als.org/stories-news/ice-bucket-challenge-dramatically-accelerated-fight-against-als

65. Samji H, Cescon A, Hogg RS, Modur SP, Althoff KN, Buchacz K, et al. Closing the Gap: Increases in Life Expectancy among Treated HIV-Positive Individuals in the United States and Canada. PLOS ONE. 2013 Dec 18;8(12):e81355.

66. Schiff D. The Importance of Facts and the Role of Academic Publishers in Today’s World—A Publisher’s View. Semin Hear. 2017 Feb;38(1):vii.

67. Nature journals raise the bar on sex and gender reporting in research. Nature. 2022 May 18;605(7910):396–396.

68. Heidari S, Babor TF, De Castro P, Tort S, Curno M. Sex and Gender Equity in Research: rationale for the SAGER guidelines and recommended use. Res Integr Peer Rev. 2016 May 3;1(1):2.

